# Evolutionary dynamics of dengue virus in India

**DOI:** 10.1101/2022.09.08.507083

**Authors:** Suraj Jagtap, Chitra Pattabiraman, Arun Sankaradoss, Sudhir Krishna, Rahul Roy

**Affiliations:** Department of Chemical Engineering, Indian Institute of Science, Bengaluru, Karnataka, India; Infectious Disease Research Foundation, Bengaluru, Karnataka, India; National Centre for Biological Sciences, Tata Institute of Fundamental Research, Bengaluru, Karnataka, India; School of Interdisciplinary Life Sciences, Indian Institute of Technology Goa, Ponda, India; Center for BioSystems Science and Engineering, Indian Institute of Science, Bengaluru, Karnataka, India

## Abstract

More than a hundred thousand dengue cases are diagnosed in India annually, and about half of the country’s population carries dengue virus-specific antibodies. Dengue propagates and adapts to the selection pressures imposed by a multitude of factors that can lead to the emergence of new variants. Yet, there has been no systematic analysis of the evolution of the dengue virus in the country. Here, we present a comprehensive analysis of all DENV gene sequences collected between 1956 and 2018 from India. We examine the spatio-temporal dynamics of India-specific genotypes, their evolutionary relationship with global and local dengue virus strains, interserotype dynamics and their divergence from the vaccine strains. Our analysis highlights the co-circulation of all DENV serotypes in India with cyclical outbreaks every 3-4 years. Since 2000, genotype III of DENV1, cosmopolitan genotype of DENV2, genotype III of DENV3 and genotype I of DENV4 have been dominating across the country. Substitution rates are comparable across the serotypes, suggesting a lack of serotype-specific evolutionary divergence. Yet, the envelope (E) protein displays strong signatures of evolution under immune selection. Apart from drifting away from its ancestors and other contemporary serotypes in general, we find evidence for recurring interserotype drift towards each other, suggesting selection via cross-reactive antibody-dependent enhancement. We identify the emergence of the highly divergent DENV4-Id lineage in South India, which has acquired half of all E gene mutations in the antigenic sites. Moreover, the DENV4-Id is drifting towards DENV1 and DENV3 clades, suggesting the role of cross-reactive antibodies in its evolution. Due to the regional restriction of the Indian genotypes and immunity-driven virus evolution in the country, ∼50% of all E gene differences with the current vaccines are focused on the antigenic sites. Our study shows how high incidence and pre-existing population immunity are shaping dengue virus evolution in India.

**Author summary:** Dengue is a mosquito-borne disease with four closely related serotypes of the virus (DENV1-4). Further, cross-reacting dengue antibodies from a previous infecting dengue serotype can protect or enhance infection from other serotypes. This can force the emergence of new dengue variants that find ways to escape the immune action or take advantage of it. In endemic countries like India, high rates of previous dengue infection can drive the evolution of dengue serotypes in complex ways. We compare all published dengue virus sequences to understand how new variants of dengue are emerging in India. Dengue cases and corresponding viruses display triennial surges. Further, the dengue envelope protein for each serotype shows recurring divergence and reversal towards its ancestral strain over a three-year window. Such fluctuations are also correlated among the dengue serotypes in India and could arise from the changing levels of cross-reactive antibodies. This, combined with the regional exchange of the virus among Asia-Pacific countries, has led to the emergence of India-specific DENV lineages, including a new DENV4 (Id) variant. This has also contributed to significant variations in the epitope regions of the current dengue viruses in India compared to the vaccines with implications for their efficacy.

## Introduction

Dengue infections have increased dramatically in the last two decades and are expected to rise further as they spread to newer regions fuelled by urbanization and travel [1]. About half of the world population is at risk of dengue infections [1,2]. Estimates from 2010 claim 390 million annual dengue infections worldwide, of which only 96 million cases were reported clinically [3]. Around one-third of these infections were estimated to be from India [3], though most of them go unreported [4]. Dengue is endemic in almost all states in India [5,6]. All four antigenically distinct serotypes (DENV1-DENV4) of the virus that display significant immunological cross-reactivity due to 65-70% homology have been reported from various parts of the country [7–9]. Combined with a complex transmission cycle and high dengue seroprevalence [5], dengue evolution in the country has been shaped in complex and unexpected ways though this remains poorly understood.

Global dengue virus evolution is modulated by pathogen transmission bottlenecks and immunological pressures [10,11]. An increase in globalization and human mobility can lead to the global spread of the emerging dengue virus strains. On the other hand, the acute nature of the disease, limited travel range and restriction to tropical regions of the vector can constrain the virus spread. Like other vector-borne virus infections, the dengue virus switches its environment due to horizontal transmission, exerting a strong purifying selection pressure [10,12,13]. In the human host as well as at the larger population scale, pre-existing immunity can contribute to the emergence of immune escape variants. Heterotypic immunity can also shape the co-evolution of dengue serotypes contingent on the level of cross-reactive antibodies and the antigenic similarity between the infecting serotypes during primary and secondary infection [11,14]. Further, antibody-dependent enhancement (ADE) under sub-optimal levels of cross-reactive antibodies can confer a selective advantage to antigenically related serotypes [15–17].

Complex population immunity against the dengue virus can also modulate the levels of annual infections and caseload. Cyclic dengue outbreaks in the endemic regions occur every 2-4 years, often associated with serotype/genotype replacement [18–21]. This has been attributed to a combination of long-term protection from homotypic dengue infection but only short-term protection (up to 2 years) from the heterotypic secondary infection [22–24]. During serotype replacement, ADE can also play a role in increasing dengue infections, depending on the level of cross-reactive immunity in the population, thereby making the cyclic pattern of outbreaks more prominent. However, whether this advantage also leads to the evolution of the virus, remains unknown. Therefore, knowledge of longitudinal prevalence, serotype distributions, and prior serotype of infection can help us in understanding the evolution of the dengue virus and predicting future outbreaks [25].

In spite of being a hotspot of dengue infections, the scarcity of dengue genomic data from India has limited our understanding of dengue virus evolution. Previous dengue studies in India have focused on regional outbreaks [7–9,26–28], which are dominated by single or closely related strains due to the limitations of a short collection period. The persistence of all serotypes in a high seroprevalence background has been shown to manifest in immunity-driven co-evolution of dengue virus strains at the city-scale [11]. In absence of longitudinal analysis of dengue viral diversity, it remains unknown, whether such selection pressures are shaping dengue virus divergence at a country-wide scale in India.

Large divergence of prevalent dengue genotypes can have significant implications for vaccine design and development [29,30]. Multiple dengue vaccines targeting all four serotypes have been developed and are currently at different stages of clinical trials [31]. These vaccines are based on the old dengue isolates from outside South Asia. In the absence of efficacy studies in India, it remains unclear whether they will induce optimal levels of neutralizing antibodies against the dengue viruses circulating in India. Apart from not providing sufficient protection against dengue infection, some vaccine candidates can even lead to enhanced disease through ADE upon subsequent infection, as seen in the case of the CYD-TDV vaccine [32,33]. Yet, differences at the antigenic sites between vaccines and prevalent Indian dengue strains have not been investigated to date.

Our group has recently published 119 whole-genome dengue sequences from clinical samples across four different sites in India from 2012 to 2018 [34]. This effort has substantially increased the number of whole-genome dengue sequences from India (from 65 to 184 genomes) that now allows careful examination of the evolutionary dynamics of the dengue virus in the country. In this study, we compiled all available whole dengue genomes (n = 184) and E gene (n = 408) sequences to generate the most comprehensive dataset of dengue virus sequences to date from India. Analysis of this dataset confirms the substantial co-circulation of DENV1, DENV2 and DENV3 in the country since 2000. Further, it shows the re-emergence of DENV4 since 2007, followed by a rapid increase in South India since 2016. The spatio-temporal analysis shows broad geographical restriction of the Indian genotypes to Asia. DENV1 genotype III, DENV2 cosmopolitan genotype (genotype IV), DENV3 genotype III and DENV4 genotype I are the dominant genotypes circulating in India. Further, the E proteins for all serotypes in India display correlated temporal fluctuations in Hamming distances with respect to its ancestors over a three-year period. Similar E gene dynamics is observed among the serotypes, pointing to the role of cross-reactive population immunity and associated ADE influencing the co-evolution of dengue serotypes. This has led to the emergence of a highly divergent DENV4 lineage in India, with evidence of strong immune selection pressure on the E gene. Dissimilarity in the vaccine genotypes and significant divergence driven by the pre-existing seropositivity also manifests in major differences in the epitopic regions compared to the vaccine strains. Our work highlights how the evolutionary dynamics of the dengue virus in India is shaped by immune selection pressure at the population level.

## Methods

### Data collection

#### Dataset A

All the published Indian dengue sequences were obtained from the ViPR database [35]. These sequences include both whole-genome sequences and gene segments. Only sequences with location and collection date were used for the analysis (about 88.9% of all sequences). Samples for these sequences were collected between 1956 and 2018 and represent DENV1 (n = 840), DENV2 (n = 877), DENV3 (n = 746) and DENV4 (n = 179) serotypes.

#### Dataset B

Global dengue protein-coding sequence records that contain sample collection dates were obtained from the ViPR database [35]. After removal of identical sequences, this dataset included DENV1 (n = 1800) from 1944-2018, DENV2 (n = 1395) from 1944-2018, DENV3 (n = 823) from 1956-2018, DENV4 (n = 220) from 1956-2018, representing a total of 4238 protein-coding sequences.

#### Subset C

For analysis of spatio-temporal dynamics, protein-coding sequences specific to the Indian genotypes were selected from dataset B. We included all unique protein-coding sequences from India and randomly sampled global sequences. This included a total of 522 sequences from DENV1 genotype I (n = 142), DENV1 genotype III (n = 93), DENV2 genotype Cosmopolitan (n = 144), DENV3 genotype III (n = 96), and DENV4 genotype I (n = 47).

#### Dataset D

All Indian E gene amino acid sequences collected since 2000 were obtained from the ViPR database [35] which comprised of DENV1 (n = 113), DENV2 (n = 168), DENV3 (n = 88), and DENV4 (n = 39) E protein sequences.

State-wise number of dengue cases and deaths over the period 2001 to 2022 was retrieved from https://www.indiastat.com/. The cases from 2019-2022 were excluded while considering the spikes due to the effect of COVID-19-related disruptions [36].

### Sequence alignment and phylogenetic analysis

Multiple sequence alignment was performed with protein-coding sequences from dataset B using MUSCLE v3.8.425 [37] implemented in AliView v1.25 [38]. Alignments were manually checked for insertion and deletion errors. Maximum likelihood trees were generated using IQ-TREE v1.6.10 [39] with 1000 bootstraps. A general time-reversible substitution model with unequal base frequency and gamma distribution for rate heterogeneity (GTR+F+I+G4) was selected out of 88 models available using jModelTest [40] implemented in IQ-TREE v1.6.10 [39] based on the Bayesian information criterion. Sylvatic strains EF457905 (DENV1), EF105379 (DENV2) and JF262779-80 (DENV4) were used as outgroups to root the respective trees. The root for DENV3 was obtained using the best fitting root by the correlation method in TempEst [41]. The maximum likelihood phylogenetic trees were used to obtain the root-to-tip distances using TempEst. Trees were visualized using Figtree v1.4. We assigned new lineages within a genotype if the difference between the sequences from each phylogenetic branch is more than 3% at the nucleotide level and 1% at the amino acid level.

Bayesian analysis was performed with the E gene, NS5 gene and whole-genome sequences from subset C using the BEAST v1.8.3 [42] to get the substitution rates for each gene. Five sets were generated for each serotype by selecting 80% sequences randomly, the substitution rate was calculated for each run, and the average substitution rate was reported. The constant rate clock model was selected based on the AICM values of the Bayes factor and harmonic means (S1 Table). Markov Chain Monte Carlo (MCMC) was run for 10^7^ generations for each run, and the first 10% of samples were discarded as burn-in. The phylogeographic movement of the virus across the countries was obtained using the Bayesian stochastic search variable selection procedure implemented in BEAST v1.8.3 with whole genome sequences from subset C (MCMC chain length ∼10^8^). SpreaD3 v0.9.7 [43] was used to visualize the spatio-temporal dynamics of the virus. Tracer v1.6 was used to check the convergence of the chains. Effective sample sizes for the parameters of interest were greater than 200.

Single-likelihood ancestral counting (SLAC) and fixed effects likelihood (FEL) methods were used to identify the positions that are undergoing selection using the HyPhy package (v2.5.1) [44]. Sites with a p-value <0.1 were considered significant only if they were detected by both methods.

### Dynamics of amino acid variation in the envelope gene sequence

Temporal dynamics of E gene amino acid variation was examined in the sequences from South India due to the availability of a larger dataset (n = 164). Sequences with at least 50% coverage (mean coverage of 93%) of the E gene were selected for the analysis. The Hamming distance between the pair of sequences was divided by the length of the overlapping region between the pair. This was further converted to the z-score using the mean and standard deviation to obtain the normalized distance. To extract the dynamics within the serotype, we measured the normalized distance of new variants over the years with respect to the ancestral sequences (sequences from 2007/08 were used as ancestors due to a lack of sufficient sequences before that). The inter-serotype distance was calculated by selecting the sequences from two different serotypes during a particular year. Sample bootstrapping (n = 100) was used to determine the median and standard error of the normalized distances for each year. The time period of oscillations was calculated using an autocorrelation function with different lags for each trace. A lag with the maximum correlation coefficient was assigned as the period of oscillation for that trace. Bootstrap replicates (n=100) were used to obtain the distribution of the time period.

Pearson’s correlation coefficient was obtained between the serotype and inter-serotype distance dynamics. The robustness of the correlations was checked by three methods. First, by randomly deleting up to 3 data points from each dynamics (1000 bootstraps). Second, by estimating the cross-correlation between individual traces from two comparing groups randomly with 1000 bootstraps. As a control, we also checked correlations between the groups by randomly shuffling the time series of normalized hamming distances (before correlating) to ensure that the correlations do not arise merely from the yearly fluctuations (1000 bootstraps).

### Dominant epitope selection from the database

Experimentally determined epitopes for all dengue serotypes were obtained from the Immunome Browser tool available on the Immune Epitope Database [45]. B cell and T cell epitopes were selected based on their response frequency (RF) score, calculated as reported earlier [46]. Epitopes having RF-score > 0.25 were selected as dominant epitopes for further analysis. For the epitopes examined in multiple studies, the RF-score was calculated by combining the number of subjects from all the studies. Apart from these known epitopes, we also included 77 sites in which residue variation conferred antigenic effects [47].

### Comparison with the vaccine strains

All Indian DENV envelope sequences post-2000 from dataset D were compared with three vaccines: CYD-TDV developed by Sanofi Pasteur, TV003 by NIH/Butantan and TAK-003 by Takeda. Multidimensional scaling based on the Hamming distances between the sequences was used to visualize and evaluate the differences between Indian and vaccine strains.

Homo-dimeric structures of the envelope proteins were obtained by homology modelling using the SWISS-MODEL server [48]. Template PDB structures were selected based on global model quality estimates and QMEAN statistics. Vaccine strain information and template selection for each serotype is shown in the S2 Table. Sites with differences in >10% of sequences were mapped onto the envelope protein structure using PyMOL (v2.4.1) [49].

## Results

### Dengue serotype dynamics in India

The reported dengue cases in 2018 have increased more than 25-fold (three year average) since 2002 in India (S1A Fig). All four geographical regions, namely– North, East, South and West-Central India, show periodic spikes in dengue cases as well as deaths over 2-4 years (S1B-D Fig). Comparing all published dengue sequences till 2018 from India (dataset A), we find all four dengue serotypes co-circulating in the country since 2000 (Fig 1). Although dengue sequence reporting from various parts of the country is sporadic, the number of annual cases and deaths in the past two decades correlated well with the number of available sequences each year (S1E Fig, Pearson’s correlation coefficient ∼ 0.65). In particular, we noted the increase in DENV2 and DENV4 sequences since 2011 and 2014, respectively (Fig 1A). Corresponding to the reported cases, we found periodic peaks in the number of sequences reported from North and South India. Genomes reported from North India show a pattern of serotype replacement in consecutive peaks (Fig 1B). Since most of the sequences from North India were collected from Delhi (∼80.5%), the spikes in the dengue sequences in 2006, 2010 and 2013 also correspond well with the outbreaks in Delhi [26,50] (S1C-D Fig). Consistent with previous reports, we find DENV3 dominated during the 2006 outbreak [51] while DENV1 and DENV2 serotypes dominated during the outbreaks in 2010 and 2013, respectively [26,52]. Although dengue outbreaks have been observed in East and West-Central India (S1C-D Fig), a relatively smaller number of sequences are available from these regions, possibly due to poor genomic surveillance (Fig 1B). Nevertheless, DENV2 emerged as the dominant serotype in East India in 2016, while all serotypes were identified in West-Central India from 2016 to 2018. In South India, three peaks are evident in the number of sequences corresponding to 2009, 2013 and 2016 (Fig 1C) and correlated peaks in the number of cases/deaths were observed in 2009-10, 2012-13 and 2017 (S1C-D Fig). DENV1 and DENV2 have been the dominant serotypes over the last decade, but DENV4 has recently emerged as a major serotype in South India. Similar periodic fluctuations in dengue incidence and replacement of dengue serotypes have been proposed previously to be driven by population-level immunity [18,19], but whether high endemicity and seropositivity are driving case incidences, serotype-specific dynamics and virus evolution in India is not known. Overall, in India, DENV1 and DENV3 were the dominating serotypes till 2012. DENV2 has become the dominant serotype in most regions in India since then, and DENV4 is establishing itself in South India.

**Fig 1.**
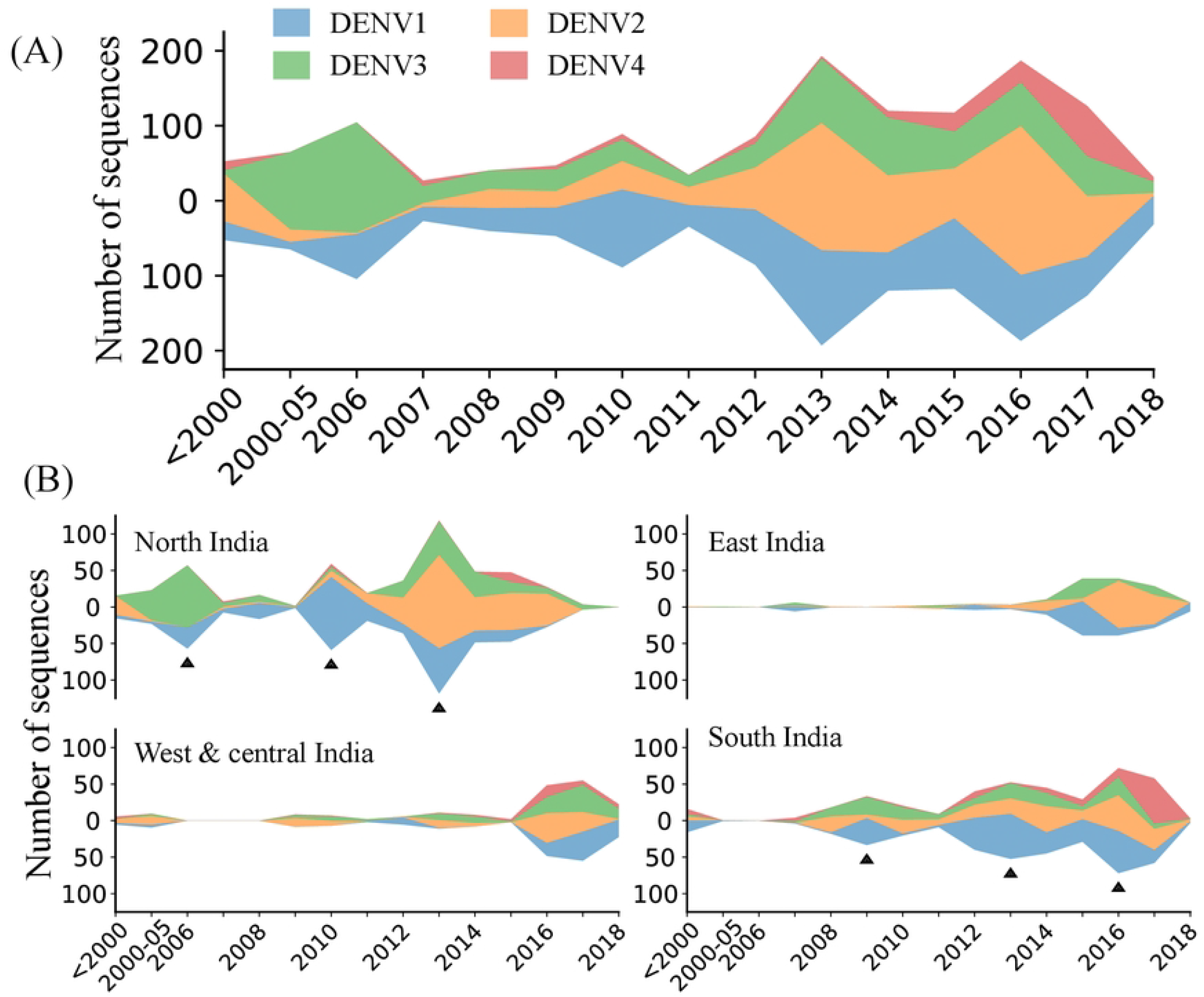
Year-wise dengue sequences reported from India. (A) Dengue serotype sequences available from India till 2018. (B) Distribution of dengue serotypes in different regions of India. DENV1: blue, DENV2: orange, DENV3: green, DENV4: red. Arrows indicate the peaks in the number of sequences.

### Prevalence and spatio-temporal dynamics of Indian dengue genotypes

Maximum likelihood trees and maximum clade credibility trees of complete genome coding sequences suggest that dengue genotypes are constrained by geography exemplified by the poor intermixing of the genotypes across continents (Fig 2, S2-4 Fig). Asian sequences consist of two dominant genotypes for all the serotypes, while the recent Indian dengue sequences represent only one of those genotypes (except for DENV1). Genotype III for DENV1, cosmopolitan genotype for DENV2, genotype III for DENV3, and genotype I for DENV4 have been the dominating genotypes in the past two decades in India. The Indian genotypes are remarkably divergent from dengue in other regions of the world beyond Asia and merit further examination of their evolutionary dynamics (Fig 2A).

**Fig 2.**
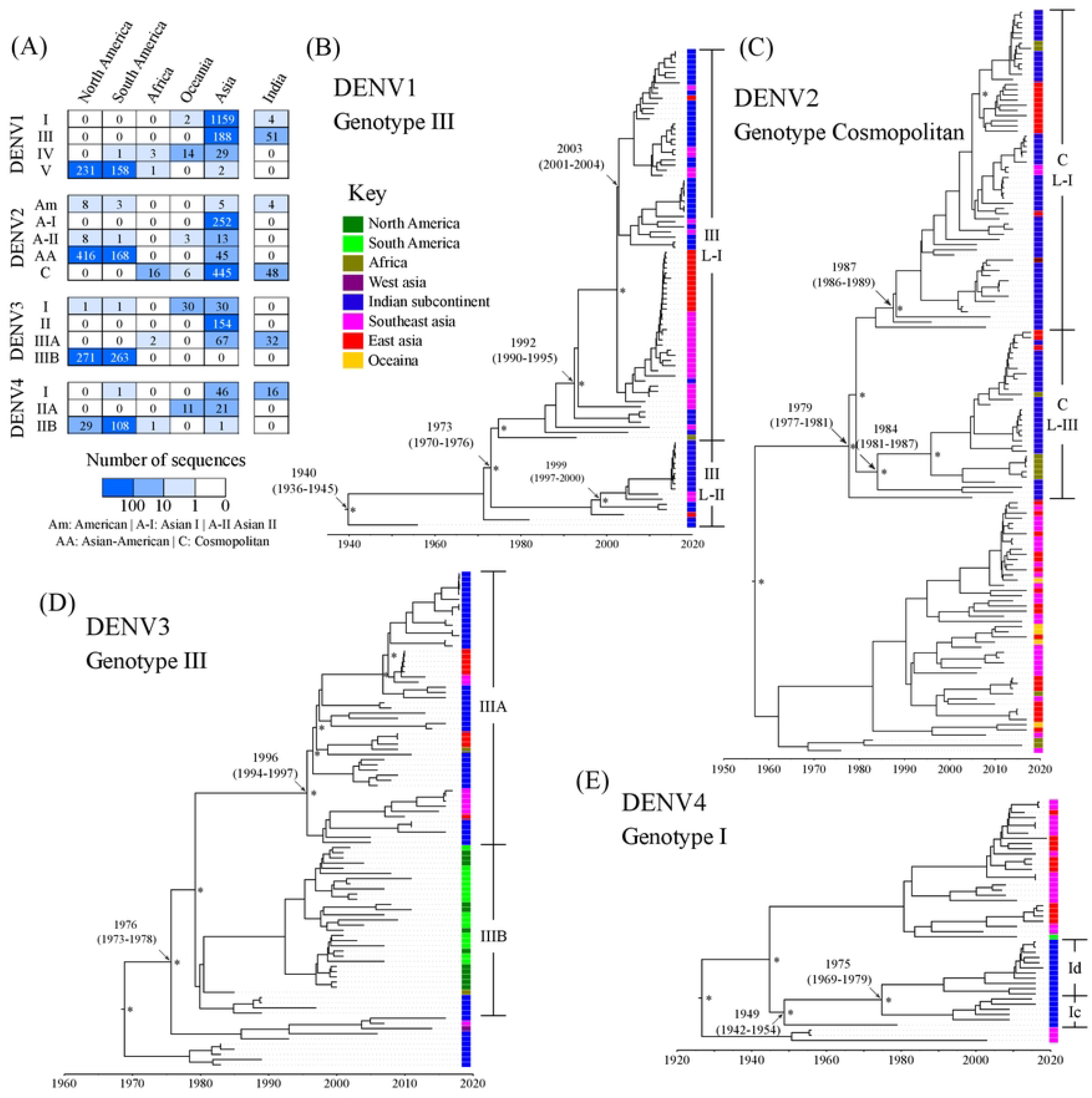
Dengue genotype distribution and phylogeny. (A) Heatmap of the number of whole-genome sequences from different geographical regions and India. (B-E) Time-dated phylogenetic trees for all serotypes with the circulating genotypes in India (DENV1-I, DENV2-cosmopolitan, DENV3-III, DENV4-I). The colour of the rectangle at the tip of the branches represents the region of the sample collection. The estimated time of the common ancestors is denoted for the important nodes along with the 95% highest posterior density (HPD) intervals. Asterisk (*) denotes posterior probability support of ≥ 0.95.

Interestingly, we find distinct temporally and spatially co-circulating lineages within the genotypes (Fig 2B-D), yet it remains unclear what factors contribute to their co-prevalence. Among the two DENV1 genotypes (I and III), genotype III has been dominant across India (S2 Fig). Within genotype III, two dominant lineages (lineage I and II) are circulating simultaneously within the country. These lineages differ at 39 positions across the genome (S5A Fig) and five positions in the immunologically dominant E gene (T272M, I337F, T339I, V358A and I/V461I). Residues 272 and 337 are on the exposed part of the E protein dimer, and position 272 is linked to having an antigenic effect suggesting differential antigenicity. Even though the phylogenetic clustering did not include the untranslated regions, all the sequences from lineage-I carry a deletion of 21 nucleotides in the hypervariable region of the 3’UTR (nucleotide positions 10294-10314 with respect to the reference sequence NC_001477, S6 Fig). A similar deletion has been reported previously from India [53], but the significance of such a large deletion is unclear. This region is not involved in any stem-loop structures or the genome cyclization region in the 3’UTR. *In vitro* studies have shown that a 19-nucleotide deletion (nucleotide positions 10289-10309) overlapping this region of 3’UTR did not affect the growth kinetics of the virus [54], while larger deletions (nucleotide positions 10274-10728) in the 3’UTR variable region show growth defects in human cell lines [55]. Therefore, it is likely that DENV1-III lineage-I can tolerate the 3’UTR deletion without a significant fitness cost. On the other hand, a cluster of sequences in DENV1-III lineage-II carry an insertion of two nucleotides in the hypervariable region of 3’UTR (C10274 and A10297 with respect to NC_001477, S6 Fig) and is restricted to South India. This shows how at least three sets of genotype III strains are co-circulating in the country. Spatio-temporal analysis reveals that DENV1-III lineage-I is a relatively new lineage that emerged in 1992 (95% HPD: 1990-1995) and is limited to India, Singapore, and China (Fig 2B, S3 Table, S7A Fig). Our analysis shows the early emergence of this genotype in India, with subsequent spread to South Korea, Comoros, and Singapore between 1990 and 2005 and multiple import-export events between India and Singapore post-2005 (S7A Fig). Apart from this, genotype I of DENV1, the primary genotype in most Asian countries (Fig 2A), has also been reported exclusively from South India since 2012, suggesting a more recent introduction (S8 Fig).

The earliest DENV2 sequences from India belong to the American genotype. This genotype was predominant in India before 1971 but was eventually replaced by the cosmopolitan genotype [56]. The DENV2-cosmopolitan genotype was introduced to India and China in the early 1980s. From here, it has spread to Southeast Asia, Australia, and East Africa (S7B Fig). DENV2 in India has two distinct co-circulating lineages (lineage-I and III) that emerged in the 80s with a common ancestor that dates back to 1979 (1977-1981) (Fig 2C, S3 Table and S3 Fig) [27,57]. While prevalent in the Indian subcontinent, cosmopolitan lineage-III has been reported in East Africa, and spatio-temporal analysis suggests that this lineage was most likely exported from India, in line with the study from Kenya [58]. These lineages differ at 41 positions across the genome, including three positions in the envelope gene (S5B Fig), all of which are in known epitopic regions or have antigenic effects (E protein residues 141, 162 and 322), suggesting immune selection pressure playing a role in their divergence. Otherwise, the E gene did not show significantly higher nonsynonymous mutation density compared to other genes (S5E Fig). Similarly, these lineages do not differ in their *in-vitro* virus growth kinetics and disease severity [27]. This could explain how these two lineages are able to avoid replacement by the other since their emergence.

All the Indian DENV3 genotype III sequences cluster along with other Asian countries (S4 Fig). DENV3-III was first detected in Sri Lanka and disseminated to multiple countries over time (S7D Fig). After its initial spread, it evolved into two distinct geographical clusters: lineages IIIA (Asian) and IIIB (American). Most of the spread of genotype IIIB in the Americas was from 1990 to 2005 (S7D Fig). In contrast, diffusion of genotype IIIA in Asia took place after 2005, yet these two lineages have not intermixed (S7D Fig, Fig 2A). In the E gene, these lineages differ at positions 158 and 380, with position 158 being a part of a known B cell epitope. Within lineage IIIA, we found two dominating sub-lineages co-circulating in India (IIIAa and IIIAb). These sub-lineages differ at 26 positions across the genome and eight positions in the E gene (S5C Fig). Although the E gene displays a high density of mutations (S5E Fig), most of these positions (7 of 8) carry the same amino acid in majority of sequences for both the sub-lineages (S5C Fig).

The spatio-temporal reconstruction identifies the origin of DENV4-I in the Philippines (S7C Fig), consistent with Sang *et al*. [59]. Most of the subsequent spread of DENV4-I was restricted to Southeast Asian countries. Although it was introduced in India around 1945, there has been a significant increase in reported DENV4-I sequences from 2015 onwards, especially in South India (Fig 1) [28,60]. It is possible that it remained undetected due to underreporting of mild infections since the primary cases of DENV4 have been reported to cause mild disease [61,62]. All Indian DENV4-I sequences cluster separately from the other prevalent genotype I groups (Fig 2E). We identify two lineages in Indian DENV4 sequences (Ic and Id); DENV4-Id lineage seems to be replacing DENV4-Ic since 2016 (Fig 2E, 3C). This novel lineage of DENV4 (Id) was reported in Pune in 2016 using the CprM sequences [63]. We noticed remarkable differences in the E and NS2A genes between the two lineages (S5D Fig), as discussed in detail in the next section. Overall, all prevalent DENV genotype lineages carry signatures of immune selection contributing to their divergence.

### Emergence of DENV4-Id suggests dominant immune selection pressure

To examine the selection pressure on the dengue virus in India, we employed FEL (Fixed Effect Likelihood) and SLAC (Single Likelihood Ancestral Counting) methods. Apart from DENV3, there was no substantial evidence of positive selection, and most amino acid changes were deleterious and negatively selected (S5, S6 Table). In DENV3-III genotype, we found a single position in NS5 protein (50I/T) under positive selection, while this site is highly conserved in all other serotypes. Significant negative selection in DENV is consistent with the requirement for horizontal transmission between taxonomically diverse host species, which imposes a strong purifying immune pressure [10,12,13].

Further, the substitution rates of dengue genotypes in India are comparable (7.59E-4, 6.31E-4, 7.83E-4, 6.50E-4 and 6.26E-4 substitutions/site/year for DENV1-I, DENV1-III, DENV2-cosmopolitan, DENV3-III and DENV4-I, respectively) (S3 Table) and similar to earlier reported rates [7,57,64]. Although 95% highest posterior density (HPD) intervals for all the genotypes overlap, the largest substitution rate is observed for the DENV2-cosmopolitan genotype. However, the substitution rates obtained for the E gene were about 26% (11.5% to 44.2%) higher than those for the whole genome (Fig 3A). Interestingly, the substitution rate was 44% larger for the DENV4-I E gene compared to the whole genome, suggesting high immunological pressure driving the divergence of the DENV4 E gene. This is consistent with previous reports for the HIV envelope gene and spike protein of the SARS-CoV-2 virus [65,66].

**Fig 3.**
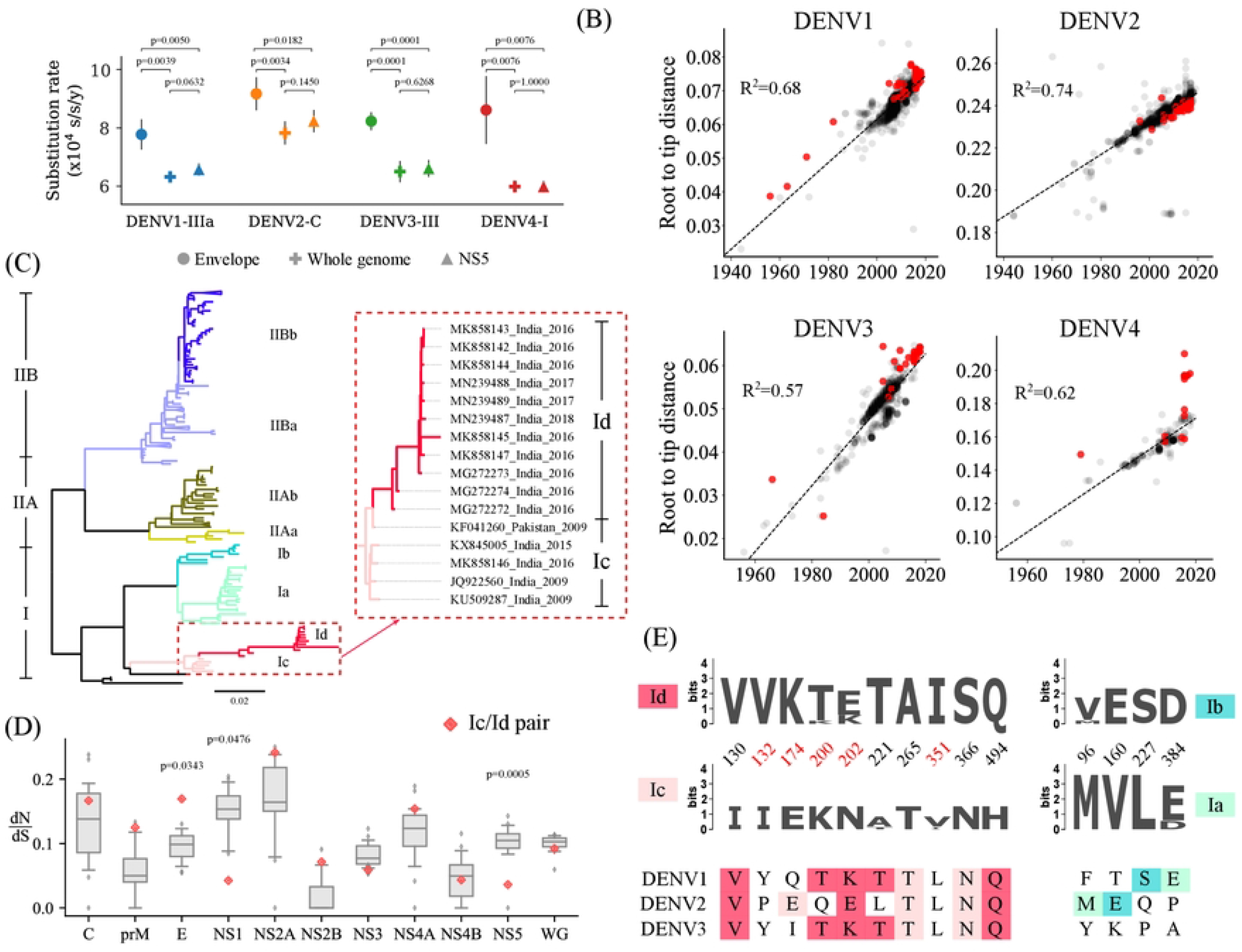
Substitution rates for Indian genotypes and DENV4-I lineage. (A) Comparison of the substitution rates (substitutions per site per year) for different segments of dengue virus for Indian genotypes. The p-values were obtained using Welch’s t-test (unpaired, two-tailed, unequal variance). (B) Root-to-tip-distance of global dengue whole-genome nucleotide sequences. The Indian sequences are highlighted in red, and the R^2^ value for the linear fit is shown. (C) The maximum-likelihood phylogenetic tree for complete coding sequences of DENV4. Distinct branches (corresponding to Ia/b/c/d, IIAa/b and IIBa/b) used for the dN/dS analysis are colour coded. Indian DENV4 cluster is shown in the inset. (D) dN/dS values for the consensus sequences generated from the branches in (C) are represented as a box plot for each gene. Whiskers denote the 5^th^ and 95^th^ percentile. The dN/dS values for Ic/d pair are marked in red. The generalized extreme Studentized deviate test was performed to find the outliers, and p-values are reported. (E) Sequence logo showing the amino acid variations between pairs Ic/d and Ia/b. The amino acids at these locations for DENV1-3 are shown at the bottom. Highlighted colour depicts the DENV4 branch with the same amino acid residue.

Root-to-tip distance analysis showed that most Indian dengue viruses follow the molecular clock similar to that observed in other parts of the world (Fig 3B). However, Indian DENV4 lineage Id is highly divergent with residuals larger than three times the interquartile range and displays the most extended branch in the phylogenetic tree (Fig 3B-C). Indeed, when we compare dN/dS ratios between branches within the DENV4 phylogenetic tree (Fig 3C), we found significantly higher dN/dS values for the E gene for the DENV4-Ic/d pair compared to other DENV4 clade pairs (Fig 3D). Besides the E gene, NS2A also showed a relatively higher dN/dS ratio and nonsynonymous mutation density (Fig 3D and S5D-E Fig). This suggests a role of immune selection pressure in DENV4-Id evolution consistent with co-evolution of E and NS2A genes correlating with virus antigenicity [47]. On the other hand, NS1 and NS5 genes showed significantly lower dN/dS ratios for the same pair. In fact, 10 out of 34 mutations in DENV4-Id (with respect to DENV4-Ic) are present in the E gene (S5D Fig). The E gene variant positions 132, 174, 200, 202 and 351 are also in the known epitopic regions and/or have antigenic effects pointing to the immune escape-driven divergence of DENV4-Id. Intriguingly, five of the acquired E gene mutations (I130V, K200T, N202K/E, A221T, H494Q) in DENV4-Id are similar to the DENV1 and DENV3 E genes (Fig 3E). This was unexpected since the prior prevalence of DENV1 and DENV3 in the country should have forced the DENV4 to drift away from them. On the other hand, antibody-dependent enhancement may confer some fitness advantage to the DENV4-Id clade due to shared antigenic features with DENV1 and DENV3. Such movement towards other serotypes is not evident in the phylogenetically related DENV4-Ia/b pair (Fig 3E), suggesting a unique signature of the Indian DENV4-Id lineage.

### Dynamics of E gene evolution displays recurring variation

Taking a cue from the divergence of DENV4-Id, we examined whether the high seroprevalence can play a role in the evolution of dengue in India. Dengue cross-reactive immunity has been shown to shape the antigenic evolution of dengue for the E gene at a city level [11]. Since serotype replacements in the different regions of the country display temporal fluctuations (Fig 1C), we asked whether there are signatures of immunity-driven evolution of dengue serotypes across large endemic areas. In absence of antigenicity data, we evaluated the variation in amino acid sequences of the E gene longitudinally. In South India, we find that, in general, the E gene diverges from the ancestral sequence for all serotypes, but this divergence fluctuates over time (Fig 4A). Overall, in our dataset, the E gene sequences drift away from their respective ancestral sequences, evolve to be similar and then diverge repeatedly. This behaviour was pronounced in DENV2-4 with an estimated time period (peak-to-peak) of about three years (2.92 ± 0.58 years for DENV2, 3.55 ± 0.9 years for DENV3 and 2.99 ± 0.64 years for DENV4). While we could not estimate a time period for a similar E gene dynamic in the case of DENV1, we did note a peak in divergence in 2012-13 arising from a singular genotype I outbreak during a period of genotype III prevalence.

**Fig 4.**
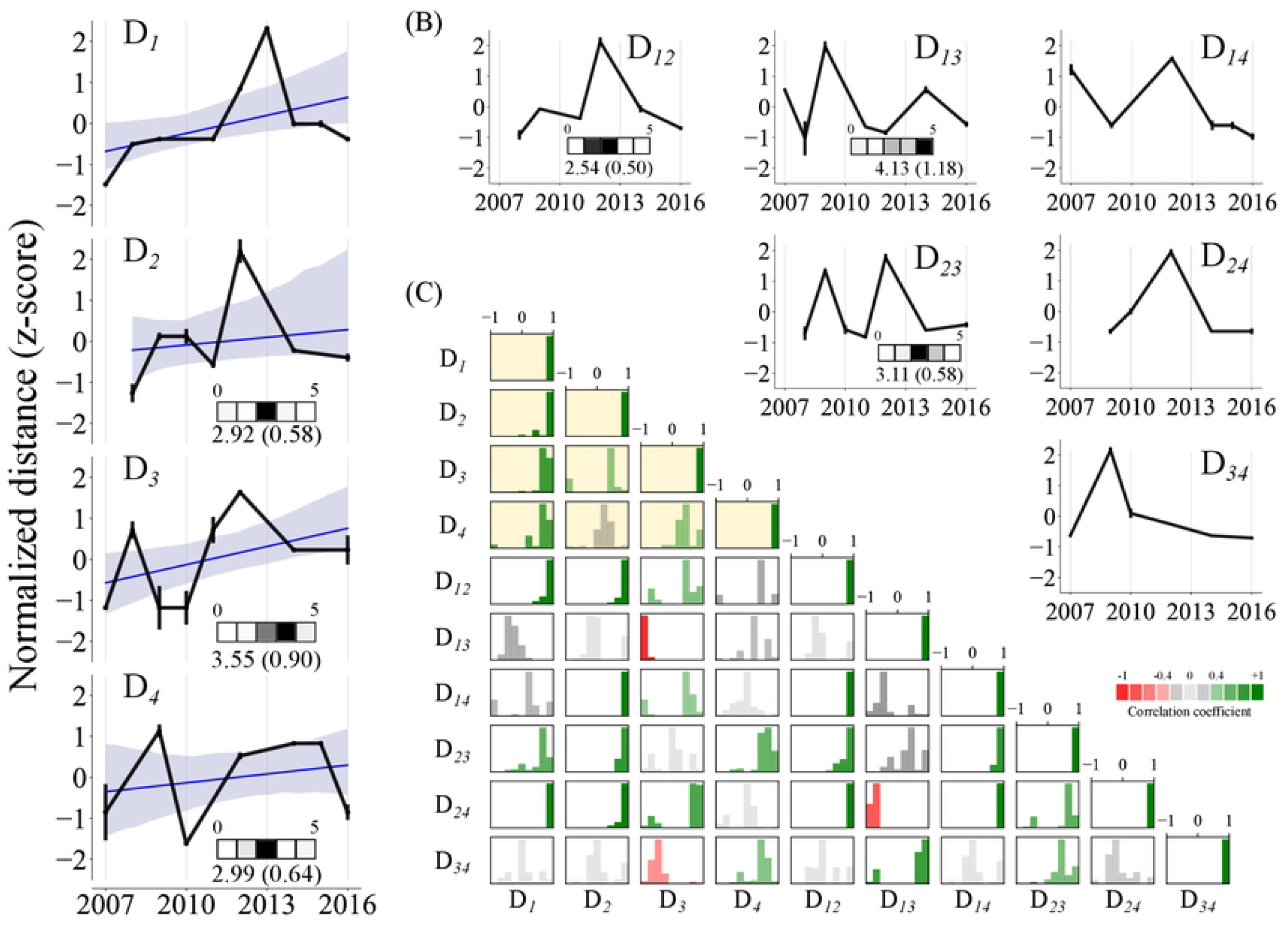
Dynamics of E gene amino acid variation in South India between 2007 and 2016. (A) Normalized amino acid distances from the strains isolated in 2007/2008. Hamming distances were normalized with the sequence length and converted to a z-score. Distances were bootstrap sampled (n = 100) to calculate the reported median. Error bars represent standard error. The blue line represents a linear regression fit to the data with 80% confidence interval(B) Year-wise normalised interserotype distances (as z-score) are reported as calculated by randomly selecting the sequences from each serotype every year (100 bootstraps). The inset heatmap depicts the time period of oscillation binned yearly (range: 0 to 5 years). The mean and standard deviation of the time period distribution is denoted next to the heatmap. (C) Distribution of correlation coefficients for within-serotype dynamics (highlighted in yellow) and inter-serotype distance dynamics. Histograms were obtained by deleting up to 3 data points randomly (1000 bootstraps). D_*i*_ indicates the distance dynamics between DENVi and its ancestral sequence. D_*ij*_ indicates the interserotype distance dynamics between *i*-th and *j*-th serotype. Green and red indicate positive (ρ > 0.4) and negative correlation (ρ < -0.4), respectively, while grey indicates weak or no correlation.

Interestingly, the observed dynamics of the E gene amino acid variations were also correlated among the serotypes (Fig 4C). The high correlation suggests that the divergence in each serotype was synchronous, i.e., when DENV1 drift away (or converges) from its ancestral sequences, a similar dynamics is observed for DENV2-4 with respect to their ancestral sequences (Fig 4C). To understand whether the evolution of the dengue virus is shaped by the interserotype cross-reactive immunity in the population, we examined the dynamics of interserotype distance *D*_*ij*_ (year-wise hamming distance between DENV*i* and DENV*j*) (Fig 4B). We observe that the interserotype distances also displayed fluctuations over a similar time period of 2-4 years (2.54 ± 0.5 years for D_*12*_, 4.13 ± 1.18 years for D_*13*_ and 3.11 ± 0.58 years for D_*23*_), suggesting an interplay between the serotypes at the population level. We also checked whether the serotype sequences became similar to each other or drifted apart based on the correlations between the serotype and interserotype evolutionary dynamics. For instance, when DENV1 and DENV2 display divergence (or convergence) with respect to their ancestral sequences (Fig 4A), the distance between the DENV1 and DENV2 (D_*12*_) also increase (or decrease). While DENV1 dynamics did not correlate with D_*13*_ and D_*14*_ dynamics, DENV2 strongly correlated with all other interserotypic dynamics. In contrast, DENV3 showed a negative correlation with D_*13*_ and D_*34*_ dynamics suggesting that as DENV3 diverges, it moves closer to the circulating DENV1 and DENV4 strains (Fig 4C). These signatures are specific to the immunologically dominant E gene. For example, when we compare the CprM gene sequences from South India, the fluctuations were not significant in any case apart from D_*1*_ dynamics but that is linked to the introduction of genotype I in 2012 (S10A Fig). Consistent with this, we found no significant correlations between the within or inter-serotypic viral dynamics, except for DENV4 (S10B-C Fig). Due to the availability of only 21 sequences for DENV4 (and n=3 between 2008-2015), we do not consider these correlations as significant.

We interpret these coupled fluctuations between dengue serotypes in light of population-level cross-reactive immunity and antibody-dependent enhancement. When homotypic immunity is present in the population, the serotypes drift apart, manifesting in positive correlations between serotype divergence and interserotype dynamics, as observed for DENV1 and DENV2. However, when population immunity is poor against a particular serotype (as with a new introduction or waning levels), similarity to that serotype confers an advantage due to the presence of cross-reactive antibodies and associated ADE [14,15]. This is consistent with the negative correlation observed between DENV3 and interserotype dynamics D_*13*_ and D_*34*_ with dropping DENV3 prevalence and its replacement by DENV1 and DENV4 in South India (Fig 1C). Indeed, recent reports have argued the interserotypic convergence of dengue antigenicity to be correlated to the outbreaks [11].

### Divergence of Indian dengue virus from vaccine strains

All the vaccine strains used by tetravalent vaccines are based on the strains isolated between 1964 and 1988. As the prevalent Indian dengue viruses are evolving under high population seropositivity, they might be diverging antigenically from the vaccine strains as well. Additionally, genotypes used in the vaccine strains are not observed in India (S2 Table). Therefore, we examined the E protein differences between the vaccine and the circulating dengue virus in India. Dimensionality reduction analysis of amino acid differences between the virus strains shows that dengue E gene sequences from India cluster together but are removed from the vaccine strains for all serotypes (Fig 5A). Only the DENV1 and DENV2 strains of CYD-TDV are close to the DENV1-I and DENV2-cosmopolitan clusters present in India. Mapping of dominant amino acid differences (present in >10% sequences) on the envelope structures shows that ≥50% of the differences (50% DENV1, 68% DENV2, 92% DENV3 and 50% DENV4) lie on the exposed surface of the envelope protein on the virus, which is accessible to the majority of the antibodies (Fig 5B, S11 Fig, S6 Table). Overall, ∼ 6% (2.7-13.5%) of all known epitopic regions are different in Indian dengue sequences compared to the vaccine strains (S7 Table). Further, almost half (34.6-66.7%) of all the E protein variations lie either in known epitopic regions or have a positive antigenic effect. This is in line with our assertion that the dengue virus in India has evolved under immunological selection pressures. It also implies that current vaccines will likely evoke poor neutralization and display limited efficacy against dengue viruses circulating in India.

**Fig 5.**
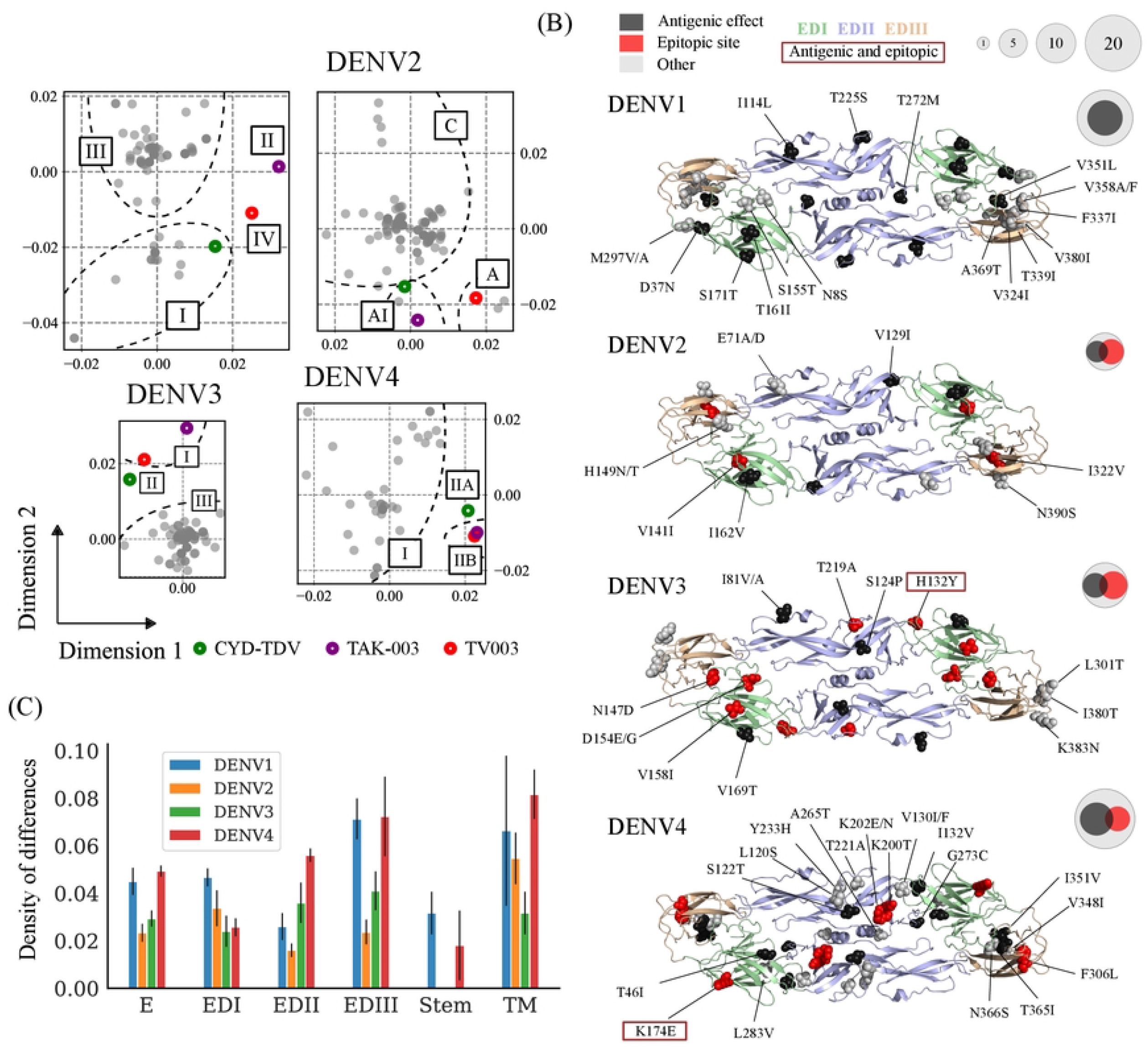
Comparison of Indian envelope protein sequences with vaccine strains. (A) Relative amino acid differences between Indian envelope sequences (post-2000) and vaccine strains (CYD-TDV, TV003 and TAK-003). The multidimensional scaling was used to reduce the dimensions of the data using the hamming distance matrix. Genotype cluster boundaries are indicated with dotted lines for visual clarity. Indian sequences are shown in grey; vaccine strains are shown with empty circles. (B) Amino acid differences at a frequency >10% with respect to the CYD-TDV vaccine are shown on the envelope protein dimer structure. Different positions in the known epitopic regions are shown in red. Variable residues within the predicted antigenic effect positions are shown in black. Residues present in epitopic regions with predicted antigenic effects are highlighted with a box. Venn diagram shows the total number of differences in the full E protein (including the stem and transmembrane region), epitopic sites and antigenic sites. The circle size represents the number of differences category-wise. (C) The density of differences (number of differences/ length of the domain) is shown for different domains of the envelope protein (envelope domains EDI-III, stem and transmembrane (TM) region). The error bars represent the standard deviation across the three vaccine strains.

Indian variants of DENV1 and DENV4 are distinct from all the vaccines compared to DENV2 and DENV3 (Fig 5C). Among the E protein domains, EDIII, a known antigenic domain and target of recent Indian dengue vaccine efforts [34,67], carry the largest density of variations, further confirming the immune escape-driven evolution of the protein. Apart from EDIII, the transmembrane region of the vaccines (especially for TAK-003) is distinctly different from the Indian variants (S6 Table). This region has been reported to be highly antigenic in DENV2 [68,69]. Consistent with the role of the stem region in the viral membrane fusion process, it is highly conserved. Therefore, antigenic differences of Indian dengue viruses with respect to the vaccines define the majority of E protein differences. This can have important implications for vaccine efficacy. Monoclonal antibodies targeting the EDIII region of dengue display a 100-fold variation in neutralization titer across different genotypes [70,71]. Similarly, challenge studies (in humans, ClinicalTrials.gov Identifier: NCT03416036 and Cynomolgus Macaques) show complete protection against vaccine genotypes but confer only partial protection against other genotypes [72,73]. For the DENV4-II based CYD-TDV vaccine, efficacy in young children against DENV4-I is only 23.9%, which has been attributed to eight specific residues in the E/prM gene [33]. Four of these (T46I, L120S, F461L, and T478S) dramatically reduce vaccine efficacy (from as high as ∼75% to less than 20%) [33]. DENV4 in TV003 and TAK-003 vaccines also share three of these residue variations, implying potentially lower vaccine efficacy against the DENV4-I genotype prevalent in India.

## Discussion

Due to the immense public health burden from dengue infections in India, it is important to understand the genetic diversity, spatial incidence, effectiveness of vaccines and the potential emergence of new variants of the dengue virus in the region. It is expected that a combination of high seroprevalence and co-circulation of all dengue serotypes shapes the evolution of the dengue virus in the country, but the outcome of the interplay between direct and indirect factors has been unclear. Consistent with immune evasion, we find that the highly immunogenic dengue E gene gradually diverges for all serotypes from their ancestral sequences over time. However, these E gene differences within each serotype were superimposed with recurrent fluctuations with a period of about three years. It is possible that opposing evolutionary constraints imparted by immune selection on one hand and E protein functional fitness on the other could manifest such fluctuations. Similar fluctuations in dengue antigenicity have been reported from Bangkok, Thailand [11], suggesting that immune selection pressure is a prominent driver for such fluctuations during virus divergence in endemic regions.

Interestingly, dengue virus interserotypic evolution is also intricately coupled to each other and displays temporally correlated fluctuations. We argue that the correlation between the intraserotype and interserotype evolutionary dynamics arises from the interplay of multiple serotypes with the pre-existing heterotypic immunity levels in India (Fig 6). Among the serotypes, antigenically related serotypes display larger cross-reactivity. The level of pre-existing cross-reactive antibodies can modulate the viral load and severity of the disease during the secondary infection [15,16]. Despite long-term protection from homotypic secondary infection, protection from reinfection with the other serotypes is transient (Fig 6A). As this cross-reactive immunity wanes, the chance of ADE increases at intermediate levels of the antibodies [15]. With further reduction in antibody titers, antibody-mediated enhancement or protection no longer contributes to the risk of infection. Therefore, high antibody titers confer protection, but intermediate levels can increase viral load and severity [15,16]. For closely related serotypes, more cross-reactive antibodies can lead to a broader ADE window without effective neutralization (Fig 6B). This can contribute to a larger viral load, longer duration of infection, higher risk of transmission and increased severity. Thus, pre-existing immunity can provide an evolutionary advantage to an antigenically ‘similar’ virus and promote convergence towards related serotypes. This could contribute to the co-evolution of ‘antigenic cousins’ as detected in our analysis of Indian dengue genotypes. When this selection pressure no longer constrains the virus (e.g., due to a reduction in cross-reactive antibodies), it is free to diverge away from the ancestral strains. Comparable time scales for observed E gene variations and the decay dynamics of interserotypic cross-reactive antibodies [22–24] suggest that such an interplay between the immune selection of dengue serotypes might be at play.

**Fig 6.**
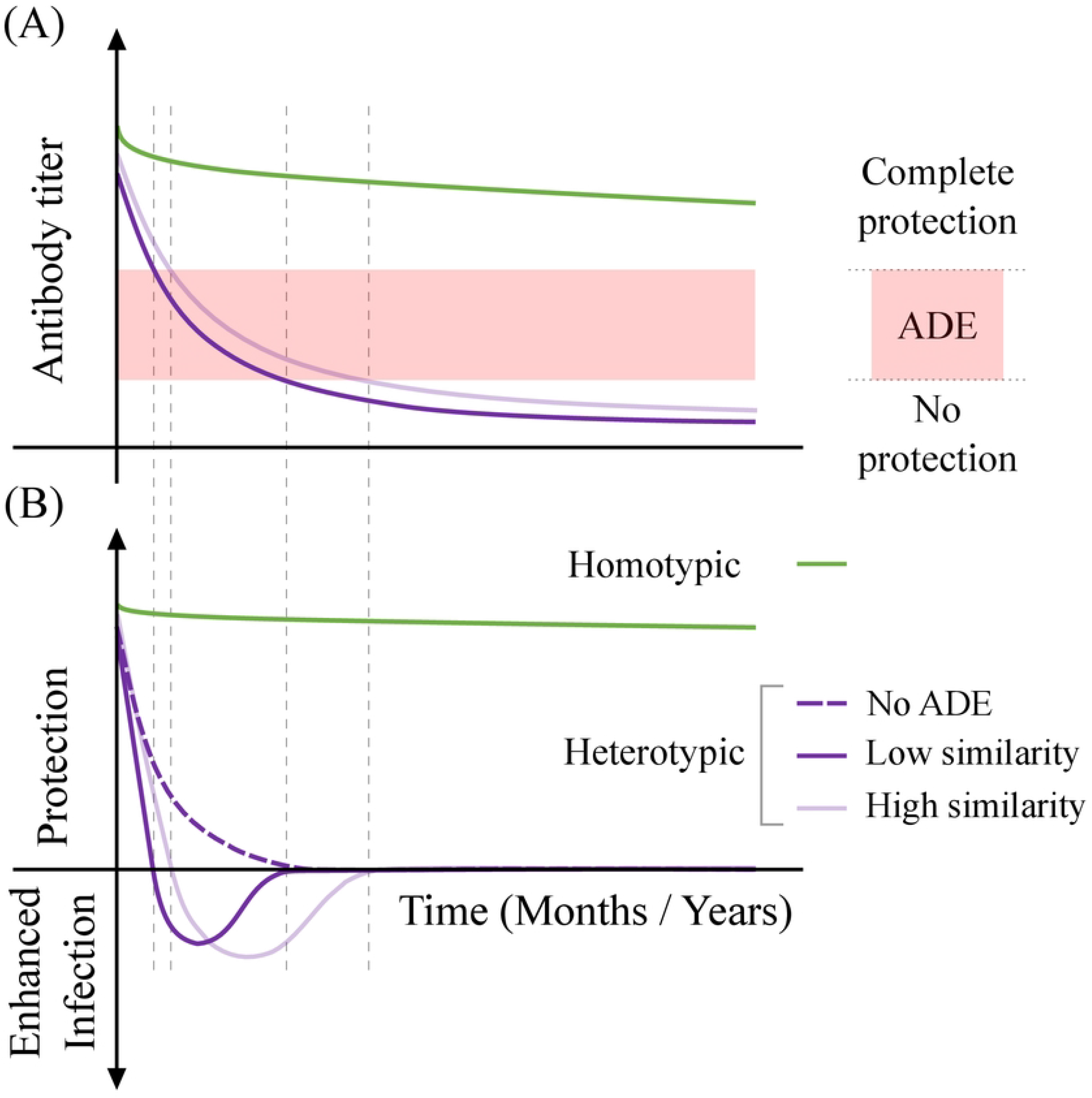
Schematic for antibody-mediated protection from secondary dengue infection. (A) Antibody decay dynamics and (B) corresponding level of protection during the secondary infection. Homotypic antibodies and protection from homotypic infection wane slowly (green). Cross-reactive antibodies to heterotypic dengue and protection wane faster (purple). Sequence similarity between the primary and secondary infecting serotypes is shown by shades of purple. The red shaded area represents the window of the antibody titer in which ADE is in effect. The dotted purple line represents protection response in the absence of ADE.

Another implication of waning protection against heterotypic infection and robust homotypic immunity contributes to serotype replacement events [19–21]. We speculate that the serotype with the cross-reactive antibody levels that manifest in strong ADE effects would have a distinctive advantage in secondary infections. This can explain how major outbreaks coincide with serotype or lineage shifts [20,21]. It also explains the close match between cross-reactive protection decay and the time interval between the heterotypic outbreak cycles in India (Fig 1, S1 Fig). Understanding the interplay of population-level immunity with dengue virus evolutionary dynamics may allow the prediction of future outbreak serotypes and genotypes.

DENV4 is emerging as one of the dominant serotypes in South India (Fig 1B) [28,60]. High divergence in the DENV4-Id lineage in India is largely restricted to immunodominant E and NS2A proteins, again pointing to immune evasion driving its emergence. Consistent with our observation of interserotype correlations, this lineage is moving towards DENV1 and DENV3 serotypes, and this enhanced similarity might contribute to ADE during the secondary infection in the Indian population. Although DENV4 was traditionally considered less virulent, it has previously replaced circulating serotypes and caused epidemics in the Pacific region[74,75]. More interestingly, a strong association of DENV4 with secondary infections (97%) observed in Thailand suggests that DENV4 can co-opt immunity-driven factors to overcome fitness limitations [76]. High seroprevalence and identification of the fast evolving ‘antigenically related’ DENV4 genotype in South India augurs an increased frequency of DENV4 outbreaks in other parts of the country and possible association with increased severity.

While the evolution of the dengue virus in India is driven by pre-existing immunity, phylogenetics also highlights the geographical restriction of dengue viral diversity (Fig 2, S7 Fig). Despite globalization and increased human mobility, dengue genotypes are contained within broad geographical boundaries, albeit with prominent intermixing between neighbouring countries [77]. This could arise from the limited lifecycle of the virus in the host and vector, climate and biodiversity. As a consequence, Indian genotypes are highly divergent from the genotypes in other continents as well as the current vaccine strains. Moreover, most of the variations from vaccine strains are concentrated on the exposed regions of the E protein, including the EDIII region, which plays a significant role in defining the antigenicity [78] and is the primary target of the neutralizing antibodies [79,80]. Therefore, an in-depth characterization of the effect of the reported differences on the antibody titers is required to assess the efficacy of prospective dengue vaccines for India. One can argue that regionally tailored vaccines would be more efficacious due to the geographical restriction of dengue [77]. However, our evolutionary analysis suggests that the immunity developed by vaccination can also impact the future course of dengue virus evolution. Further, the evolution of dengue serotypes (and other flaviviruses [81]) mediated by heterotypic immunity argues that their dynamics is intertwined that cannot be inferred fully by studying them separately. As the incidences of dengue virus continue to increase worldwide, the host immunity-driven virus evolution would need to be considered carefully to devise interventions.

## Acknowledgements

We would like to thank Mary Dias, Guruprasad Madigeshi, Jayanthi Shastri, Ravisekhar Gadepalli, and Saranya Marimuthu for their help and valuable feedback on this work.

